# Weed species composition and diversity of 2-year-old oil palm tree/fruit vegetables intercrop in rainforest zone of Nigeria

**DOI:** 10.1101/2020.05.31.126722

**Authors:** Ayodele Samuel Oluwatobi

**Affiliations:** Crown-Hill University, Faculty of Science, Department of Natural and Environmental Sciences, Eiyenkorin, Ilorin, Nigeria

**Keywords:** crop, diversity, immature, intercropping, oil palm, weed

## Abstract

The spacing pattern and growth habit of juvenile oil palm during the early stages of field establishment have often led to serious weed problem until canopy closure at subsequent years. This study was carried out during the rainy season of 2016 to evaluate the weed species composition and diversity of an intercrop between 2-year-old oil palm tree and two fruit vegetables at an oil palm plantation in Ala, Akure-North Local Government, Ondo state, Nigeria. Two accessions of tomato (NGB 01665 and NG/AA/SEP/09/053) and eggplant (NGB 01737) were intercropped within the alley of immature oil palm. Weed sampling was carried out using 0.25 m^2^ quadrat within each experimental plot. Quantitative analysis of weed species parameters and Simpson’s Diversity Index were evaluated. The result revealed that 28, 21 and 20 weed species were found across all the plots at 3, 6 weeks after intercropping (WAI), and after harvesting respectively. Members of Asteraceae produced the highest weed species at 3 WAI (17.857%); Poaceae recorded the highest weed species at 6 WAI and after harvesting (19.048% and 20%) respectively. A total of 23, 16 and 15 broadleaves were found at 3 and 6 WAI, and after harvesting. In all the juvenile oil palm/vegetable intercrops evaluated, the control plot recorded the highest weed species richness at 6 WAI, when compared to other intercropping regimes. The control and juvenile oil palm/tomato (NGB 01665) intercrop plot recorded the highest and lowest Simpson’s Indices of Diversity at 6 WAI (0.877 and 0.734) respectively.

## Introduction

The wide field spacing (6 – 9 m between palm stands) used in oil palm cultivation allows for waste of solar emission and has consequently led to serious weed infestation from transplanting to canopy closure stages when the oil palm becomes productive. Oil palms possess a long juvenility phase of about 4 years during which the palm is not productive. During this period, farmers spend on a lot of money on labour for weeding the plantation every year without any monetary returns (Oluwatobi, 2019).

The broad empty alley among the rows of immature oil palms have culminated in inefficient or ineffective employment of environmental resources, growing space, land, CO2 and sunlight (Rezig *et al*, 2012). Thus, annual crops introduction, like soybean and groundnut in juvenile oil palm plantation areas, has furnished with the opportunity to enhance the effective and efficient utilization of resources in the environment (Rezig *et al*, 2012; Abera and Feyisa, 2009; Aynehband *et al*, 2010; Amanullah *et al*, 2006).

Weed management is one of the most important economic and agronomic issues facing farmers (Rew *et al.*, 2005). Evaluating the impacts of different intercropping system and component crop combinations on weed distribution is important in selecting more efficient intercropping patterns and crop combinations that will assist to lessen weed infestation greatly, develop better weed management systems, increase yield; and hence promote food security. Unmindful of the cropping schemes employed, the weed problem remains a major cause of yield loss in crops and detailed knowledge and understanding of their distribution, biology, survival mechanism and life cycle would help in further research to reduce distressing effects of weeds on agricultural farms and ultimately food security. Weed species distribution in terms of species richness is measured as the number of species in a community. Distribution could be within or between communities; two communities with the same number of species can differ in terms of evenness, and hence it is useful to know the proportion or relative abundance of species within the community (Karaye *et al*., 2007).

Therefore, the aim of this study was to evaluate the weed species composition and distribution in juvenile oil palm/fruit vegetables intercrop. The objectives were to evaluate the weed species composition in intercropped alley and control plots; weed species richness and abundance; and weed species diversity.

## Materials and Methods

### Study area

The field experiment was conducted within an established juvenile oil palm plantation located at Ala in Akure-North Local Government Area of Ondo State. The oil palm plantation is located at a coordinate range of Latitude 7.093° N, Longitude 5.354°E (N7°5’ 35.59837” E5°21’ 15.47179”), and Latitude 7.09302 Longitude 5.35422 (N7°5’ 34.8857” E5°21’ 15.19177”) in the tropical rain forest region of Nigeria. It has two distinct seasons namely: dry and rainy seasons. Rainy season is between April and November and dry season is between November and March. Annual rainfall varies from 1150 to 2550 mm. Temperature is moderately high year-round and range between 22°C and 34°C with daily average of 30°C (Ogunrayi *et al.,* 2016).

### Weed Collection

Weed survey was carried out according to the method of Olorunmaiye *et al.* (2011). An ‘M’ pattern was systematically identified and mapped out in each plot from the edge and 0.5 × 0.5 m quadrat (0.25m^2^) was used for sampling 5 times per plot (for area less than 5 hectares).

Weeds found within each quadrat were uprooted, sorted into species, identified, counted and recorded. Data collected were subjected to ecological analysis according to Oluwatobi and Olorunmaiye (2014). Weeds found within each quadrat in each plot were identified using a standard Handbook of West African Weeds by Akobundu and Agyakwa (1998). They were counted and recorded to compute the frequency, density, relative frequency (RF), relative density (RD) and relative importance value (RIV), abundance, dominance, and relative dominance of each species according to Das (2011) and Oluwatobi and Olorunmaiye (2014).

Prior to each weeding at 3 and 6 weeks after intercropping (WAI), weed species were collected to determine weed density (counts m^−2^) and biomass (grams m^−2^). Weeds were cut at ground level, packed in papers bags, oven dried at 80°C for 2 hours and weighed to determine their biomass (Ogunkunle *et* al., 2013; Chipomho *et al.*, 2015).

### Weed species parameters and diversity

Data collected were computed to determine quantitative analysis such as frequency, density, relative frequency, relative density, importance value, abundance, dominance and relative dominance. These parameters were determined according to Olorunmaiye *et al.* (2013) and Oluwatobi and Olorunmaiye (2014). Frequency of weeds was computed as number of quadrats where weed species was found. Weed density was estimated as number of species per area of quadrat (0.25 m^2^). Other parameters were determined as follows:

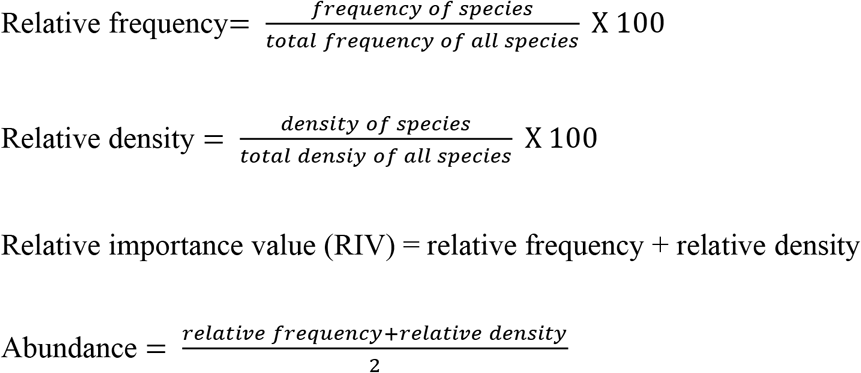

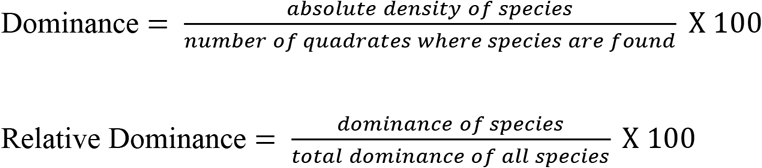

Simpson’s Diversity Index (D): This estimates the likelihood that two individual weed species erratically selected from a sample will fit in to the same species. This index was calculated as follows:

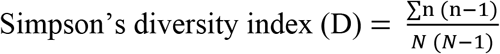

Where n = total number of a particular weed species; N = total number of all weed species

Simpson’s index of diversity (1 – D): This is the likelihood that two individual weed species arbitrarily selected from a sample will belong to different species. This was calculated by subtracting Simpson’s diversity index (D) from 1.

Simpson’s reciprocal index (1/D): The value of this index commences with 1 as the least likely figure. The figure would signify a community comprising only one species. It was calculated by finding the inverse of Simpson’s index of diversity (D) (Ali and Bibi, 2013; Oluwatobi and Olorunmaiye, 2014).

Weed smothering efficiency was determined using the mathematical formula below:

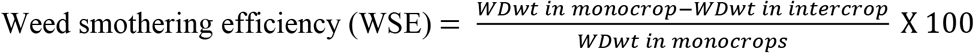

WD_wt_ = weed dry weight. (Chipomho et al., 2015).

The data gathered from the study were analyzed statistically using statistical package for social sciences (SPSS: version 17.0) by subjecting the data to analysis of variance (ANOVA). The mean values of the data were separated using Duncan Multiple Range Test (DMRT) at 5% probability level. The results were presented in tables.

## Results

### Weed Species Composition

Twenty-eight (28) different weed species were found at 3 weeks after intercropping (WAI) across the 7 plots sampled. Five species belong to Asteraceae (17.857%), 4 species belong to each of Euphorbiaceae and Poaceae, 2 belong to Malvaceae, 1 species belongs to each of Rubiaceae, Commelinaceae, Convolvulaceae, Cyperaceae, Cleomaceae, Lamiaceae, Portulaceae, Leguminosae-Papilionoideae, Urticaceae, Aizoaceae, Piperaceae, Nyctaginaceae and Loganiaceae.

At 6 WAI, 21 weed species were found across the 7 plots sampled. Four (4) of the weed species belong to Poaceae family (19.048%); 3 weed species each belong to families Asteraceae and Euphorbiaceae; 1 weed species each belong to families Portulaceae, Commelinaceae, Cyperaceae, Piperaceae, Rubiaceae, Solanaceae, Malvaceae, Loganiaceae, Nyctagianaceae, Lamiaceae and Cleomaceae.

After consecutive harvesting of all the fruits from the vegetables, 20 weed species were found across the 7 plots sampled. Four (4) weed species belong to Poaceae family (20%); 3 weed species belong to Euphorbiaceae family; 2 weed species each belong to Asteraceae and Solanaceae; and 1 weed species each belongs to Commelinaceae, Cyperaceae, Rubiaceae, Loganiaceae, Piperaceae, Malvaceae, Nyctaginaceae, Portulaceae and Leguminosae-papilionoideae.

The result showed a reduction in the number of weed species at 3 WAI (28) through 6 WAI (21) till after harvesting of fruits from the intercropped vegetables (20).

Summarily:

At 3 WAI, a total of 23 broadleaves, 4 grasses and 1 sedge weed species (82.14, 14.29 and 3.57%) respectively were found.
At 6 WAI, a total of 16 broadleaves, 4 grasses and 1 sedge weed species (76.19, 19.05 and 4.76%) respectively were found.

After harvesting, a total of 15 broadleaves, 4 grasses and 1 sedge weed species (75.00, 20.00 and 5.00%) were found. Results of weed flora and species parameters at fruit vegetable/ juvenile oil palm intercropped plots are presented in Tables 1 – 12.

**Table 1.**
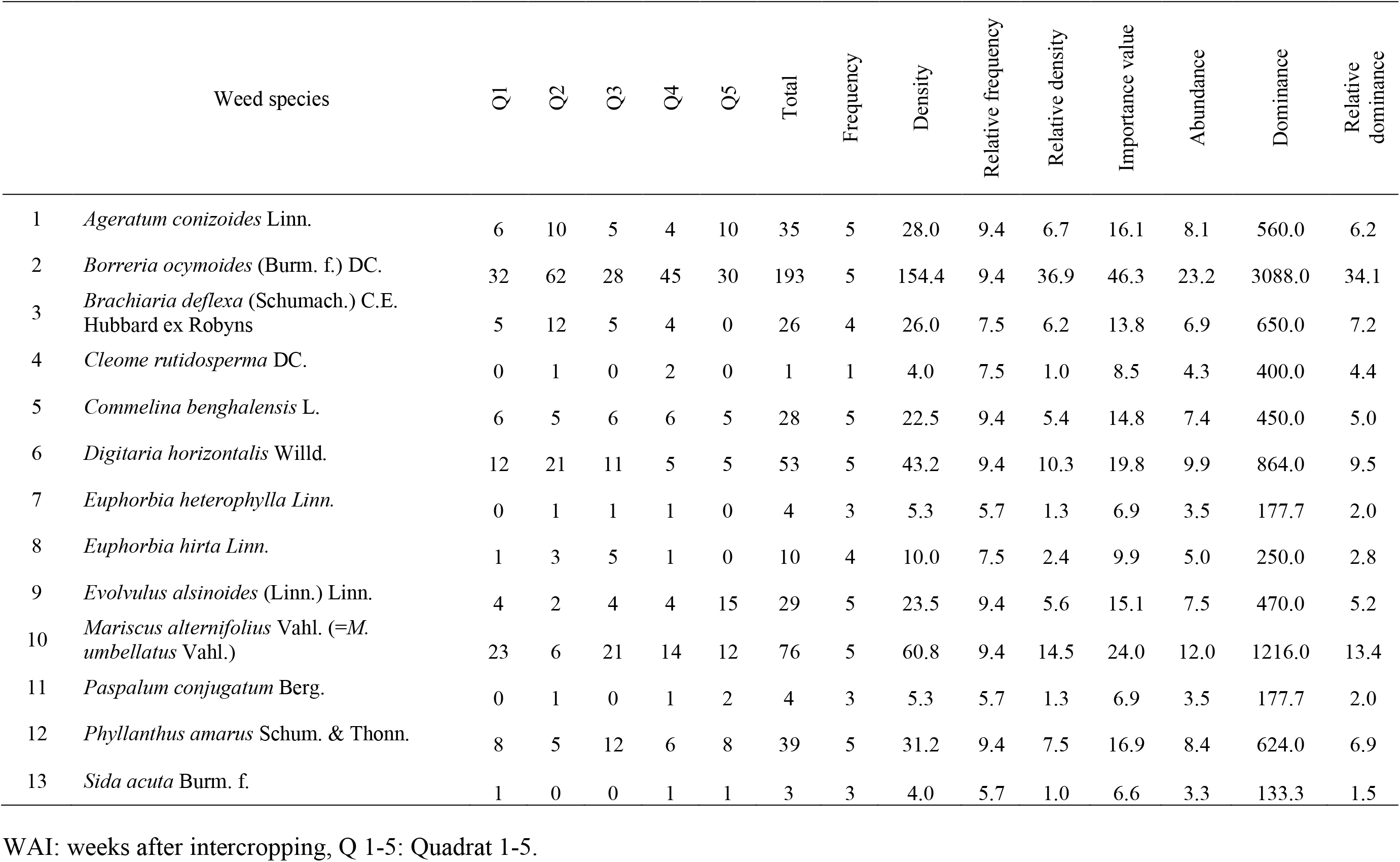
Weed flora and species parameters for tomato (NGB 01665) plot at 3 WAI

|  | Weed species | Q1 | Q2 | Q3 | Q4 | Q5 | Total | Frequency | Density | Relative frequency | Relative density | Importance value | Abundance | Dominance | Relative dominance |
| --- | --- | --- | --- | --- | --- | --- | --- | --- | --- | --- | --- | --- | --- | --- | --- |
| 1 | *Ageratum conizoides* Linn. | 6 | 10 | 5 | 4 | 10 | 35 | 5 | 28.0 | 9.4 | 6.7 | 16.1 | 8.1 | 560.0 | 6.2 |
| 2 | *Borreria ocymoides* (Burm. f.) DC. | 32 | 62 | 28 | 45 | 30 | 193 | 5 | 154.4 | 9.4 | 36.9 | 46.3 | 23.2 | 3088.0 | 34.1 |
| 3 | *Brachiaria deflexa* (Schumach.) C.E. Hubbard ex Robyns | 5 | 12 | 5 | 4 | 0 | 26 | 4 | 26.0 | 7.5 | 6.2 | 13.8 | 6.9 | 650.0 | 7.2 |
| 4 | *Cleome rutidosperma* DC. | 0 | 1 | 0 | 2 | 0 | 1 | 1 | 4.0 | 7.5 | 1.0 | 8.5 | 4.3 | 400.0 | 4.4 |
| 5 | *Commelina benghalensis* L. | 6 | 5 | 6 | 6 | 5 | 28 | 5 | 22.5 | 9.4 | 5.4 | 14.8 | 7.4 | 450.0 | 5.0 |
| 6 | *Digitaria horizontalis* Willd. | 12 | 21 | 11 | 5 | 5 | 53 | 5 | 43.2 | 9.4 | 10.3 | 19.8 | 9.9 | 864.0 | 9.5 |
| 7 | *Euphorbia heterophylla* Linn. | 0 | 1 | 1 | 1 | 0 | 4 | 3 | 5.3 | 5.7 | 1.3 | 6.9 | 3.5 | 177.7 | 2.0 |
| 8 | *Euphorbia hirta* Linn. | 1 | 3 | 5 | 1 | 0 | 10 | 4 | 10.0 | 7.5 | 2.4 | 9.9 | 5.0 | 250.0 | 2.8 |
| 9 | *Evolvulus alsinoides* (Linn.) Linn. | 4 | 2 | 4 | 4 | 15 | 29 | 5 | 23.5 | 9.4 | 5.6 | 15.1 | 7.5 | 470.0 | 5.2 |
| 10 | *Mariscus alternifolius* Vahl. (=M. umbellatus Vahl.) | 23 | 6 | 21 | 14 | 12 | 76 | 5 | 60.8 | 9.4 | 14.5 | 24.0 | 12.0 | 1216.0 | 13.4 |
| 11 | *Paspalum conjugatum* Berg. | 0 | 1 | 0 | 1 | 2 | 4 | 3 | 5.3 | 5.7 | 1.3 | 6.9 | 3.5 | 177.7 | 2.0 |
| 12 | *Phyllanthus amarus* Schum. & Thonn. | 8 | 5 | 12 | 6 | 8 | 39 | 5 | 31.2 | 9.4 | 7.5 | 16.9 | 8.4 | 624.0 | 6.9 |
| 13 | *Sida acuta* Burm. f. | 1 | 0 | 0 | 1 | 1 | 3 | 3 | 4.0 | 5.7 | 1.0 | 6.6 | 3.3 | 133.3 | 1.5 |

WAI: weeks after intercropping, Q 1-5: Quadrat 1-5.

**Table 2.** Weed flora and species parameters for tomato (NGB 01665) plot at 6 WAI

|  | Weed species | Q1 | Q2 | Q3 | Q4 | Q5 | Total | Frequency | Density | Relative frequency | Relative density | Importance value | Abundance | Dominance | Relative dominance |
| --- | --- | --- | --- | --- | --- | --- | --- | --- | --- | --- | --- | --- | --- | --- | --- |
| 1 | *Ageratum conizoides* Linn. | 15 | 6 | 6 | 8 | 4 | 39 | 5 | 31.2 | 10.9 | 7.6 | 18.5 | 9.2 | 624.0 | 7.0 |
| 2 | *Bidens pilosa* Linn. | 6 | 13 | 0 | 7 | 5 | 31 | 4 | 31.0 | 8.7 | 7.6 | 16.3 | 8.1 | 775.0 | 8.6 |
| 3 | *Boerhavia diffusa* L. | 2 | 10 | 1 | 6 | 6 | 25 | 5 | 20.0 | 10.9 | 4.9 | 15.7 | 7.9 | 400.0 | 4.5 |
| 4 | *Borreria ocymoides* (Burm. f.) DC. | 6 | 16 | 18 | 6 | 9 | 49 | 5 | 39.2 | 10.9 | 9.6 | 20.4 | 10.2 | 784.0 | 8.7 |
| 5 | *Commelina benghalensis* L. | 1 | 1 | 0 | 0 | 0 | 2 | 2 | 4.0 | 4.3 | 1.0 | 5.3 | 2.7 | 200.0 | 2.2 |
| 6 | *Croton lobatus* L. | 2 | 2 | 0 | 1 | 0 | 5 | 3 | 6.7 | 6.5 | 1.6 | 8.1 | 4.1 | 222.2 | 2.5 |
| 7 | *Hyptis lanceolata* Poir. | 2 | 1 | 0 | 0 | 0 | 3 | 2 | 6.0 | 4.3 | 1.5 | 5.8 | 2.9 | 300.0 | 3.3 |
| 8 | *Mariscus alternifolius* Vahl. (=M. *umbellatus* Vahl.) | 15 | 10 | 9 | 15 | 6 | 55 | 5 | 44.0 | 10.9 | 10.7 | 21.6 | 10.8 | 880.0 | 9.8 |
| 9 | *Peperomia pellucida* (L.) H. B. & K. | 30 | 28 | 56 | 41 | 76 | 231 | 5 | 184.8 | 10.9 | 45.1 | 55.9 | 28.0 | 3696.0 | 41.2 |
| 10 | *Setaria barbata* (Lam.) Kunth | 18 | 6 | 5 | 3 | 7 | 39 | 5 | 31.2 | 10.9 | 7.6 | 18.5 | 9.2 | 624.0 | 7.0 |
| 11 | *Sida acuta* Burm. f. | 0 | 0 | 1 | 0 | 1 | 2 | 2 | 4.0 | 4.3 | 1.0 | 5.3 | 2.7 | 200.0 | 2.2 |
| 12 | *Solanum nigrum* L. | 3 | 2 | 0 | 1 | 0 | 6 | 3 | 8.0 | 6.5 | 2.0 | 8.5 | 4.2 | 266.7 | 3.0 |

WAI: weeks after intercropping, Q 1-5: Quadrat 1-5.

**Table 3.** Weed flora and species parameters for tomato (NGB 01665) plot after harvesting

|  | Weed species | Q1 | Q2 | Q3 | Q4 | Q5 | Total | Frequency | Density | Relative frequency | Relative density | Importance value | Abundance | Dominance | Relative dominance |
| --- | --- | --- | --- | --- | --- | --- | --- | --- | --- | --- | --- | --- | --- | --- | --- |
| 1 | *Ageratum conizoides* Linn. | 53 | 32 | 28 | 41 | 32 | 185 | 5 | 148.0 | 10.2 | 34.9 | 45.1 | 22.5 | 2960.0 | 31.3 |
| 2 | *Bidens pilosa* Linn. | 0 | 3 | 0 | 2 | 8 | 13 | 3 | 17.3 | 6.1 | 4.1 | 10.2 | 5.1 | 577.8 | 6.1 |
| 3 | *Borreria ocymoides* (Burm. f.) DC. | 3 | 4 | 12 | 9 | 15 | 43 | 5 | 34.4 | 10.2 | 8.1 | 18.3 | 9.2 | 688.0 | 7.3 |
| 4 | *Commelina benghalensis* L. | 8 | 14 | 14 | 10 | 6 | 52 | 5 | 41.6 | 10.2 | 9.8 | 20.0 | 10.0 | 832.0 | 8.8 |
| 5 | *Euphorbia heterophylla* Linn. | 0 | 4 | 0 | 4 | 3 | 11 | 3 | 14.7 | 6.1 | 3.5 | 9.6 | 4.8 | 488.9 | 5.2 |
| 5 | *Mariscus alternifolius* Vahl. (=M. umbellatus Vahl.) | 6 | 13 | 4 | 6 | 3 | 32 | 5 | 25.6 | 10.2 | 6.0 | 16.2 | 8.1 | 512.0 | 5.4 |
| 6 | *Peperomia pellucida* (L.) H. B. & K. | 0 | 2 | 5 | 0 | 8 | 15 | 3 | 20.0 | 6.1 | 4.7 | 10.8 | 5.4 | 666.7 | 7.1 |
| 7 | *Phyllanthus amarus* Schum. & Thonn. | 1 | 5 | 0 | 3 | 2 | 11 | 4 | 11.0 | 8.2 | 2.6 | 10.8 | 5.4 | 275.0 | 2.9 |
| 8 | *Physalis angulata* Linn. | 3 | 1 | 0 | 1 | 0 | 4 | 3 | 5.3 | 6.1 | 1.3 | 7.4 | 3.7 | 177.8 | 1.9 |
| 9 | *Spigelia anthelmia* Linn. | 20 | 25 | 12 | 10 | 19 | 86 | 5 | 68.8 | 10.2 | 16.2 | 26.4 | 13.2 | 1376.0 | 14.6 |
| 10 | *Solanum nigrum* L. | 2 | 10 | 2 | 14 | 6 | 34 | 5 | 27.2 | 10.2 | 6.4 | 16.6 | 8.3 | 544.0 | 5.8 |

WAI: weeks after intercropping, Q 1-5: Quadrat 1-5.

**Table 4.** Weed flora and species parameters for tomato (NG/AA/SEP/09/053) plot at 3 WAI

| Weed species |  | Q1 | Q2 | Q3 | Q4 | Q5 | Total | Frequency | Density | Relative frequency | Relative density | Importance value | Abundance | Dominance | Relative dominance |
| --- | --- | --- | --- | --- | --- | --- | --- | --- | --- | --- | --- | --- | --- | --- | --- |
| 1 | *Ageratum conizoides* Linn. | 3 | 15 | 16 | 18 | 10 | 62 | 5 | 49.6 | 9.6 | 16.5 | 26.1 | 13.0 | 992.0 | 11.9 |
| 2 | *Axonopus compressus* (Sw.) P. Beauv. | 0 | 1 | 1 | 2 | 3 | 7 | 4 | 7.0 | 7.7 | 2.3 | 10.2 | 5.0 | 175.0 | 2.1 |
| 3 | *Bidens pilosa* Linn. | 1 | 0 | 1 | 0 | 2 | 4 | 3 | 5.3 | 5.8 | 1.8 | 7.5 | 3.8 | 177.7 | 2.1 |
| 4 | *Borreria ocymoides* (Burm. f.) DC. | 38 | 5 | 6 | 5 | 8 | 62 | 5 | 49.6 | 9.6 | 16.5 | 26.1 | 13.0 | 992.0 | 11.9 |
| 5 | *Brachiaria deflexa* (Schumach.) C.E. Hubbard ex Robyns | 12 | 8 | 8 | 10 | 4 | 42 | 5 | 33.6 | 9.6 | 11.2 | 20.8 | 10.4 | 672.0 | 7.5 |
| 6 | *Digitaria horizontalis* Willd. | 7 | 9 | 3 | 9 | 10 | 38 | 5 | 30.4 | 9.6 | 10.1 | 19.7 | 9.9 | 608.0 | 7.3 |
| 7 | *Euphorbia hirta* Linn. | 0 | 1 | 0 | 0 | 0 | 1 | 1 | 4.0 | 1.9 | 2.3 | 3.3 | 1.6 | 400.0 | 4.8 |
| 8 | *Evolvulus alsinoides* (Linn.) Linn. | 0 | 2 | 0 | 3 | 1 | 6 | 3 | 8.0 | 5.8 | 2.7 | 8.4 | 4.2 | 266.7 | 3.2 |
| 9 | *Hyptis lanceolata* Poir. | 2 | 0 | 0 | 0 | 0 | 2 | 1 | 8.0 | 1.9 | 2.7 | 4.6 | 2.3 | 800.0 | 9.6 |
| 10 | *Mariscus alternifolius* Vahl. (=M. *umbellatus* Vahl.) | 2 | 0 | 1 | 0 | 0 | 3 | 2 | 6.0 | 3.8 | 2.0 | 5.8 | 2.9 | 300.0 | 3.6 |
| 11 | *Paspalum conjugatum* Berg. | 0 | 2 | 0 | 0 | 2 | 4 | 2 | 8.0 | 3.8 | 2.7 | 6.5 | 3.3 | 400.0 | 4.8 |
| 12 | *Pouzolzia guineensis* Benth. | 0 | 1 | 1 | 0 | 0 | 2 | 2 | 4.0 | 3.8 | 1.3 | 5.2 | 2.6 | 200.0 | 2.4 |
| 13 | *Synedrella nodiflora* Gaertn. | 5 | 16 | 15 | 8 | 6 | 50 | 5 | 40.0 | 9.6 | 13.3 | 22.9 | 11.4 | 800.0 | 9.6 |
| 14 | *Talinum triangulare* (Jacq.) Willd. | 0 | 4 | 4 | 0 | 6 | 14 | 3 | 18.7 | 5.8 | 6.2 | 12.0 | 6.0 | 622.3 | 7.5 |
| 15 | *Tephrosia pedicellata* Bak. | 0 | 3 | 0 | 1 | 0 | 4 | 2 | 8.0 | 3.8 | 2.7 | 6.5 | 3.3 | 400.0 | 4.8 |
| 16 | *Tridax procumbens* Linn. | 8 | 4 | 3 | 0 | 6 | 21 | 4 | 21.0 | 7.7 | 7.0 | 14.7 | 7.3 | 525.0 | 6.3 |

WAI: weeks after intercropping, Q 1-5: Quadrat 1-5.

**Table 5.** Weed flora and species parameters for tomato (NG/AA/SEP/09/053) plot at 6 WAI

|  | Weed species | Q1 | Q2 | Q3 | Q4 | Q5 | Total | Frequency | Density | Relative frequency | Relative density | Importance value | Abundance | Dominance | Relative dominance |
| --- | --- | --- | --- | --- | --- | --- | --- | --- | --- | --- | --- | --- | --- | --- | --- |
| 1 | *Ageratum conizoides* Linn. | 21 | 14 | 12 | 6 | 14 | 67 | 5 | 53.6 | 12.2 | 20.6 | 32.8 | 16.4 | 1072.0 | 17.7 |
| 2 | *Bidens pilosa* Linn. | 4 | 18 | 5 | 5 | 14 | 46 | 5 | 36.8 | 12.2 | 14.1 | 26.3 | 13.2 | 736.0 | 12.2 |
| 3 | *Borreria ocymoides* (Burm. f.) DC. | 0 | 4 | 1 | 3 | 0 | 8 | 3 | 10.7 | 7.3 | 4.1 | 11.4 | 5.7 | 355.6 | 5.9 |
| 4 | *Commelina benghalensis* L. | 17 | 2 | 10 | 10 | 6 | 45 | 5 | 36.0 | 12.2 | 13.8 | 26.0 | 13.0 | 720.0 | 11.9 |
| 5 | *Euphorbia hirta* Linn. | 0 | 2 | 1 | 0 | 2 | 5 | 3 | 6.7 | 7.3 | 2.6 | 9.9 | 4.9 | 233.3 | 3.9 |
| 6 | *Mariscus alternifolius* Vahl. (= *M. umbellatus* Vahl.) | 0 | 2 | 2 | 0 | 0 | 4 | 2 | 8.0 | 4.9 | 3.0 | 7.9 | 4.0 | 400.0 | 6.6 |
| 7 | *Peperomia pellucida* (L.) H. B. & K. | 10 | 7 | 5 | 12 | 7 | 41 | 5 | 32.8 | 12.2 | 12.6 | 24.8 | 12.4 | 656.0 | 10.8 |
| 8 | *Phyllanthus amarus* Schum. & Thonn. | 0 | 2 | 0 | 1 | 1 | 4 | 3 | 5.3 | 7.3 | 2.0 | 9.4 | 4.7 | 177.7 | 2.9 |
| 9 | *Setaria barbata* (Lam.) Kunth | 11 | 25 | 17 | 11 | 9 | 73 | 5 | 58.4 | 12.2 | 22.4 | 34.0 | 17.3 | 1168.0 | 876.0 |
| 10 | *Solanum nigrum* L. | 0 | 1 | 1 | 1 | 0 | 3 | 3 | 4.0 | 7.3 | 1.5 | 8.9 | 4.4 | 133.3 | 2.2 |
| 11 | *Talinum triangulare* (Jacq.) Willd. | 1 | 0 | 0 | 3 | 0 | 4 | 2 | 8.0 | 4.9 | 3.0 | 7.9 | 4.0 | 400.0 | 6.6 |

WAI: weeks after intercropping, Q 1-5: Quadrat 1-5.

**Table 6.** Weed flora and species parameters for tomato (NG/AA/SEP/09/053) plot after harvesting

| Weed species |  | Q1 | Q2 | Q3 | Q4 | Q5 | Total | Frequency | Density | Relative frequency | Relative density | Importance value | Abundance | Dominance | Relative dominance |
| --- | --- | --- | --- | --- | --- | --- | --- | --- | --- | --- | --- | --- | --- | --- | --- |
| 1 | *Ageratum conizoides* Linn. | 5 | 2 | 0 | 12 | 6 | 25 | 4 | 25.0 | 9.3 | 8.1 | 17.4 | 8.7 | 625.0 | 8.1 |
| 2 | *Bidens pilosa* Linn. | 1 | 0 | 1 | 2 | 2 | 6 | 4 | 6.0 | 9.3 | 2.0 | 11.6 | 5.6 | 150.0 | 1.9 |
| 3 | *Borreria ocymoides* (Burm. f.) DC. | 36 | 25 | 38 | 16 | 18 | 133 | 5 | 106.4 | 11.6 | 34.6 | 46.3 | 23.1 | 2128.0 | 27.5 |
| 4 | *Brachiaria deflexa* (Schumach.) C.E. Hubbard ex Robyns | 13 | 6 | 6 | 2 | 9 | 36 | 5 | 28.8 | 11.6 | 9.4 | 21.0 | 10.5 | 576.0 | 7.4 |
| 5 | *Brachiaria lata* (Schumach) C.E. Hubbard | 8 | 8 | 6 | 2 | 0 | 24 | 4 | 24.0 | 9.3 | 7.8 | 17.1 | 8.6 | 600.0 | 7.7 |
| 6 | *Commelina benghalensis* L. | 0 | 0 | 6 | 4 | 4 | 14 | 3 | 18.7 | 7.0 | 6.1 | 13.1 | 6.5 | 622.2 | 8.0 |
| 7 | *Euphorbia hirta* Linn. | 0 | 1 | 0 | 0 | 1 | 2 | 2 | 4.0 | 4.7 | 1.3 | 6.0 | 3.0 | 200.0 | 2.6 |
| 8 | *Mariscus alternifolius* Vahl. (=M. *umbellatus* Vahl.) | 7 | 7 | 1 | 6 | 2 | 23 | 5 | 18.4 | 11.6 | 6.0 | 17.6 | 8.8 | 368.0 | 4.7 |
| 9 | *Phyllanthus amarus* Schum. & Thonn. | 4 | 3 | 11 | 7 | 7 | 32 | 5 | 25.6 | 11.6 | 8.3 | 20.0 | 10.0 | 517.0 | 6.6 |
| 10 | *Setaria barbata* (Lam.) Kunth | 6 | 4 | 15 | 13 | 10 | 48 | 5 | 38.4 | 11.6 | 12.5 | 24.1 | 12.1 | 768.0 | 9.9 |

WAI: weeks after intercropping, Q 1-5: Quadrat 1-5.

**Table 7.** Weed flora and species parameters for eggplant (NGB 01737) plot at 3 WAI

| Weed species |  | Q1 | Q2 | Q3 | Q4 | Q5 | Total | Frequency | Density | Relative frequency | Relative density | Importance value | Abundance | Dominance | Relative dominance |
| --- | --- | --- | --- | --- | --- | --- | --- | --- | --- | --- | --- | --- | --- | --- | --- |
| 1 | *Ageratum conizoides* Linn. | 3 | 5 | 12 | 4 | 0 | 24 | 4 | 24.0 | 6.5 | 5.8 | 12.3 | 6.1 | 600.0 | 6.3 |
| 2 | *Bidens pilosa* Linn. | 1 | 0 | 0 | 0 | 2 | 3 | 2 | 6.0 | 3.2 | 1.5 | 4.7 | 2.3 | 300.0 | 3.1 |
| 3 | *Boerhavia diffusa* L. | 1 | 0 | 1 | 2 | 0 | 4 | 3 | 5.3 | 4.8 | 1.3 | 6.1 | 3.1 | 177.7 | 1.9 |
| 4 | *Borreria ocymoides* (Burm. f.) DC. | 4 | 4 | 0 | 0 | 6 | 14 | 3 | 18.7 | 4.8 | 4.5 | 9.4 | 4.7 | 622.3 | 6.5 |
| 5 | *Brachiaria deflexa* (Schumach.) C.E. Hubbard ex Robyns | 8 |  | 6 | 8 | 5 | 27 | 4 | 27.0 | 6.5 | 6.6 | 13.0 | 6.5 | 675.0 | 7.0 |
| 6 | *Cleome rutidosperma* DC. | 0 | 2 | 2 | 1 | 3 | 8 | 4 | 8.0 | 6.5 | 1.9 | 8.4 | 4.2 | 200.0 | 2.1 |
| 7 | *Commelina benghalensis* L. | 0 | 5 | 5 | 2 | 12 | 24 | 4 | 24.0 | 6.5 | 5.8 | 12.3 | 6.1 | 600.0 | 6.3 |
| 8 | *Euphorbia heterophylla* Linn. | 3 | 0 | 1 | 2 | 3 | 9 | 4 | 9.0 | 6.5 | 2.2 | 8.6 | 4.3 | 225.0 | 2.3 |
| 9 | *Euphorbia hirta* Linn. | 0 | 2 | 1 | 0 | 3 | 6 | 3 | 8.0 | 4.8 | 1.9 | 6.8 | 3.4 | 266.7 | 2.8 |
| 10 | *Mariscus alternifolius* Vahl. (=M. *umbellatus* Vahl.) | 4 | 5 | 18 | 8 | 15 | 50 | 5 | 40.0 | 8.1 | 9.7 | 17.8 | 8.9 | 800.0 | 8.3 |
| 11 | *Paspalum conjugatum* Berg. | 9 | 0 | 12 | 10 | 6 | 37 | 4 | 37.0 | 6.5 | 9.0 | 15.5 | 7.7 | 925.0 | 9.6 |
| 12 | *Peperomia pellucida* (L.) H. B. & K. | 2 | 18 | 22 | 3 | 9 | 54 | 5 | 43.2 | 8.1 | 10.5 | 18.6 | 9.3 | 864.0 | 9.0 |
| 13 | *Phyllanthus amarus* Schum. & Thonn. | 1 | 0 | 0 | 3 | 1 | 5 | 3 | 6.7 | 4.8 | 1.6 | 6.5 | 3.2 | 222.3 | 2.3 |
| 14 | *Spigelia anthelmia* Linn. | 2 | 1 | 1 | 1 | 2 | 7 | 5 | 5.6 | 8.1 | 1.4 | 9.4 | 4.7 | 112.0 | 1.2 |
| 15 | *Synedrella nodiflora* Gaertn. | 44 | 26 | 46 | 30 | 32 | 178 | 5 | 142.4 | 8.1 | 34.7 | 42.7 | 21.4 | 2848.0 | 29.7 |
| 16 | *Talinum triangulare* (Jacq.) Willd. | 0 | 2 | 1 | 2 | 1 | 6 | 4 | 6.0 | 6.5 | 1.5 | 7.9 | 4.0 | 150.0 | 1.6 |

WAI: weeks after intercropping, Q 1-5: Quadrat 1-5.

**Table 8.** Weed flora and species parameters for eggplant (NGB 01737) plot at 6 WAI

| Weed species |  | Q1 | Q2 | Q3 | Q4 | Q5 | Total | Frequency | Density | Relative frequency | Relative density | Importance value | Abundance | Dominance | Relative dominance |
| --- | --- | --- | --- | --- | --- | --- | --- | --- | --- | --- | --- | --- | --- | --- | --- |
| 1 | *Ageratum conizoides* Linn. | 25 | 22 | 28 | 20 | 18 | 113 | 5 | 90.4 | 11.1 | 27.3 | 38.4 | 19.2 | 1808.0 | 24.0 |
| 2 | *Borreria ocymoides* (Burm. f.) DC. | 12 | 14 | 10 | 15 | 28 | 79 | 5 | 63.2 | 11.1 | 19.1 | 30.2 | 15.1 | 1264.0 | 16.8 |
| 3 | *Brachiaria lata* (Schumach) C.E. | 1 | 0 | 0 | 0 | 0 | 1 | 1 | 4.0 | 2.2 | 1.2 | 3.4 | 1.7 | 400.0 | 5.3 |
| 4 | *Cleome rutidosperma* DC. | 1 | 0 | 0 | 1 | 1 | 3 | 3 | 4.0 | 6.7 | 1.2 | 7.9 | 3.9 | 133.3 | 1.8 |
| 5 | *Commelina benghalensis* L. | 5 | 6 | 5 | 5 | 8 | 29 | 5 | 23.2 | 11.1 | 7.0 | 18.1 | 9.1 | 464.0 | 6.2 |
| 6 | *Mariscus alternifolius* Vahl. (=M. *umbellatus* Vahl.) | 1 | 1 | 0 | 1 | 1 | 4 | 4 | 4.0 | 8.9 | 1.2 | 10.1 | 5.0 | 100.0 | 1.3 |
| 7 | *Peperomia pellucida* (L.) H. B. & K. | 7 | 3 | 2 | 9 | 4 | 25 | 5 | 20.0 | 11.1 | 6.0 | 17.1 | 8.6 | 400.0 | 5.3 |
| 8 | *Phyllanthus amarus* Schum. & Thonn. | 0 | 3 | 1 | 2 | 0 | 6 | 3 | 8.0 | 6.7 | 2.4 | 9.1 | 4.5 | 266.7 | 3.5 |
| 9 | *Setaria barbata* (Lam.) Kunth | 16 | 31 | 41 | 10 | 12 | 110 | 5 | 88.0 | 11.1 | 26.5 | 37.7 | 18.8 | 1760.0 | 23.4 |
| 10 | *Spigelia anthelmia* Linn. | 3 | 6 | 2 | 1 | 0 | 12 | 4 | 12.0 | 8.9 | 3.6 | 12.5 | 6.3 | 300.0 | 4.0 |
| 11 | *Talinum triangulare* (Jacq.) Willd. | 1 | 3 | 1 | 0 | 0 | 5 | 3 | 6.7 | 6.7 | 2.0 | 8.7 | 4.3 | 222.2 | 3.0 |
| 12 | *Tridax procumbens* Linn. | 3 | 1 | 0 | 0 | 0 | 4 | 2 | 8.0 | 4.4 | 2.4 | 6.9 | 3.4 | 400.0 | 5.3 |

WAI: weeks after intercropping, Q 1-5: Quadrat 1-5.

**Table 9.**
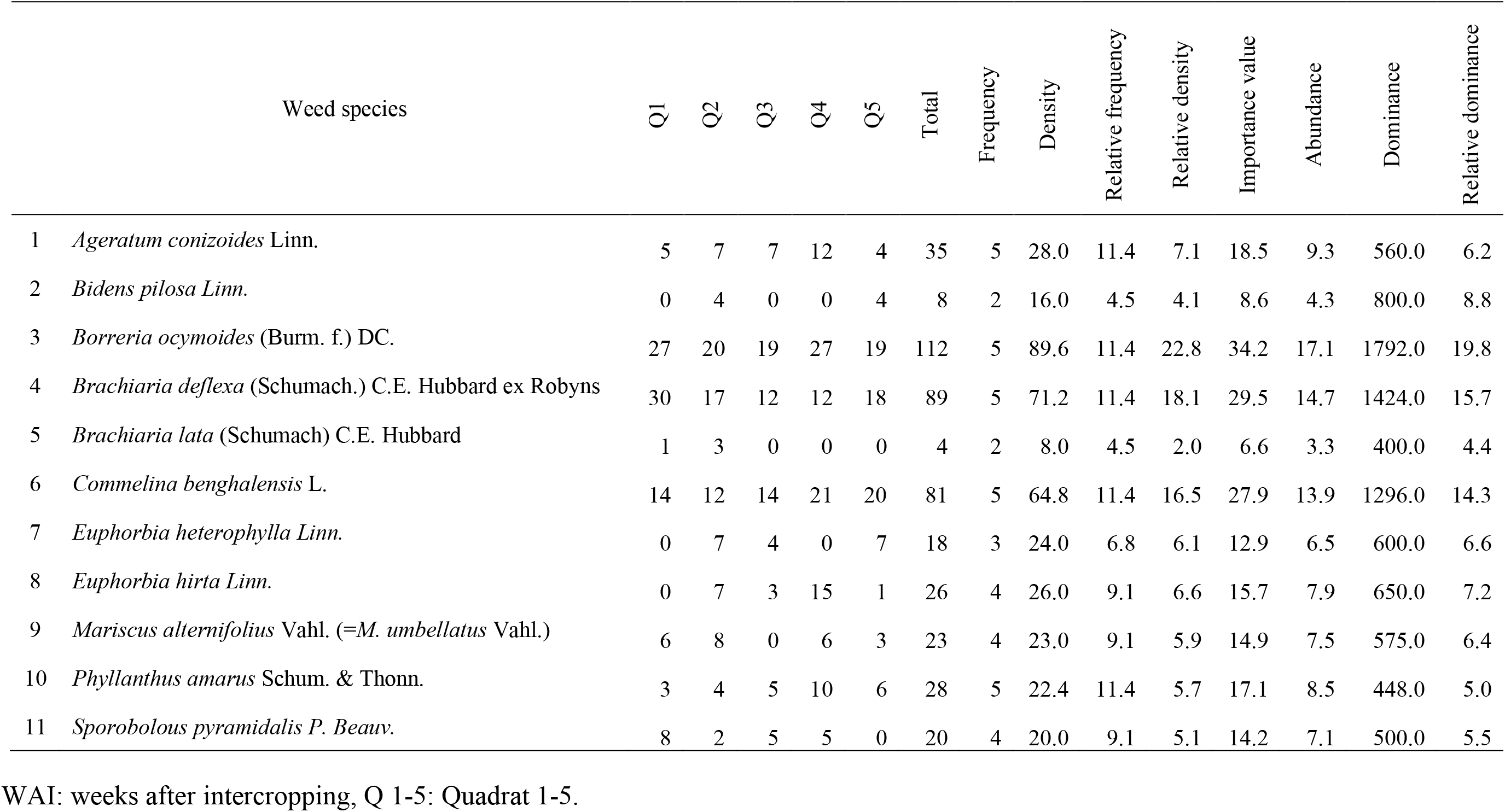
Weed flora and species parameters for eggplant (NGB 01737) plot after harvesting

| Weed species |  | Q1 | Q2 | Q3 | Q4 | Q5 | Total | Frequency | Density | Relative frequency | Relative density | Importance value | Abundance | Dominance | Relative dominance |
| --- | --- | --- | --- | --- | --- | --- | --- | --- | --- | --- | --- | --- | --- | --- | --- |
| 1 | *Ageratum conizoides* Linn. | 5 | 7 | 7 | 12 | 4 | 35 | 5 | 28.0 | 11.4 | 7.1 | 18.5 | 9.3 | 560.0 | 6.2 |
| 2 | *Bidens pilosa* Linn. | 0 | 4 | 0 | 0 | 4 | 8 | 2 | 16.0 | 4.5 | 4.1 | 8.6 | 4.3 | 800.0 | 8.8 |
| 3 | *Borreria ocymoides* (Burm. f.) DC. | 27 | 20 | 19 | 27 | 19 | 112 | 5 | 89.6 | 11.4 | 22.8 | 34.2 | 17.1 | 1792.0 | 19.8 |
| 4 | *Brachiaria deflexa* (Schumach.) C.E. Hubbard ex Robyns | 30 | 17 | 12 | 12 | 18 | 89 | 5 | 71.2 | 11.4 | 18.1 | 29.5 | 14.7 | 1424.0 | 15.7 |
| 5 | *Brachiaria lata* (Schumach) C.E. Hubbard | 1 | 3 | 0 | 0 | 0 | 4 | 2 | 8.0 | 4.5 | 2.0 | 6.6 | 3.3 | 400.0 | 4.4 |
| 6 | *Commelina benghalensis* L. | 14 | 12 | 14 | 21 | 20 | 81 | 5 | 64.8 | 11.4 | 16.5 | 27.9 | 13.9 | 1296.0 | 14.3 |
| 7 | *Euphorbia heterophylla* Linn. | 0 | 7 | 4 | 0 | 7 | 18 | 3 | 24.0 | 6.8 | 6.1 | 12.9 | 6.5 | 600.0 | 6.6 |
| 8 | *Euphorbia hirta* Linn. | 0 | 7 | 3 | 15 | 1 | 26 | 4 | 26.0 | 9.1 | 6.6 | 15.7 | 7.9 | 650.0 | 7.2 |
| 9 | *Mariscus alternifolius* Vahl. (=M. *umbellatus* Vahl.) | 6 | 8 | 0 | 6 | 3 | 23 | 4 | 23.0 | 9.1 | 5.9 | 14.9 | 7.5 | 575.0 | 6.4 |
| 10 | *Phyllanthus amarus* Schum. & Thonn. | 3 | 4 | 5 | 10 | 6 | 28 | 5 | 22.4 | 11.4 | 5.7 | 17.1 | 8.5 | 448.0 | 5.0 |
| 11 | *Sporobolous pyramidalis* P. Beauv. | 8 | 2 | 5 | 5 | 0 | 20 | 4 | 20.0 | 9.1 | 5.1 | 14.2 | 7.1 | 500.0 | 5.5 |

WAI: weeks after intercropping, Q 1-5: Quadrat 1-5.

**Table 10.** Weed flora and species parameters for control plot at 3 WAI

| Weed species |  | Q1 | Q2 | Q3 | Q4 | Q5 | Total | Frequency | Density | Relative frequency | Relative density | Importance value | Abundance | Dominance | Relative dominance |
| --- | --- | --- | --- | --- | --- | --- | --- | --- | --- | --- | --- | --- | --- | --- | --- |
| 1 | *Ageratum conizoides* Linn. | 81 | 106 | 78 | 64 | 30 | 269 | 5 | 53.8 | 7.7 | 16.1 | 23.8 | 11.9 | 1076.0 | 15.1 |
| 2 | *Bidens pilosa* Linn. | 12 | 18 | 6 | 12 | 4 | 52 | 5 | 41.6 | 7.7 | 12.4 | 20.1 | 10.1 | 832.0 | 11.7 |
| 3 | *Borreria ocymoides* (Burm. f.) DC. | 6 | 24 | 42 | 6 | 7 | 85 | 5 | 68.0 | 7.7 | 20.3 | 28.0 | 14.0 | 1360.0 | 19.1 |
| 4 | *Brachiaria deflexa* (Schumach.) C.E. Hubbard ex Robyns | 12 | 10 | 8 | 14 | 3 | 47 | 5 | 37.6 | 7.7 | 11.2 | 18.9 | 9.5 | 752.0 | 10.6 |
| 5 | *Cleome rutidosperma* DC. | 1 | 6 | 3 | 1 | 1 | 12 | 5 | 9.6 | 7.7 | 2.9 | 10.6 | 5.3 | 192.0 | 2.7 |
| 6 | *Commelina benghalensis* L. | 2 | 1 | 0 | 1 | 0 | 4 | 3 | 5.3 | 4.6 | 1.6 | 6.2 | 3.1 | 177.7 | 2.5 |
| 7 | *Digitaria horizontalis* Willd. | 8 | 12 | 9 | 3 | 1 | 33 | 5 | 26.4 | 7.7 | 7.9 | 15.6 | 7.8 | 528.0 | 7.4 |
| 8 | *Euphorbia hirta* Linn. | 1 | 5 | 0 | 2 | 1 | 9 | 4 | 9.0 | 6.2 | 2.7 | 8.8 | 4.4 | 225.0 | 3.2 |
| 9 | *Mariscus alternifolius* Vahl. (= *M. umbellatus* Vahl.) | 5 | 5 | 2 | 11 | 6 | 29 | 5 | 23.2 | 7.7 | 6.9 | 14.6 | 7.3 | 464.0 | 6.5 |
| 10 | *Peperomia pellucida* (L.) H. B. & K. | 3 | 19 | 4 | 2 | 6 | 34 | 5 | 27.2 | 7.7 | 8.1 | 15.8 | 7.9 | 544.0 | 7.6 |
| 11 | *Spigelia anthelmia* Linn. | 1 | 4 | 0 | 2 | 0 | 7 | 3 | 9.3 | 4.6 | 2.8 | 7.4 | 3.7 | 311.0 | 4.4 |
| 12 | *Synedrella nodiflora* Gaertn. | 1 | 6 | 5 | 0 | 2 | 14 | 4 | 14.0 | 6.2 | 4.2 | 10.3 | 5.2 | 350.0 | 4.9 |
| 13 | *Talinum triangulare* (Jacq.) Willd. | 1 | 4 | 0 | 0 | 2 | 7 | 3 | 9.3 | 4.6 | 2.8 | 7.4 | 3.7 | 311.0 | 4.4 |

WAI: weeks after intercropping, Q 1-5: Quadrat 1-5.

**Table 11.** Weed flora and species parameters for control plot at 6 WAI

| Weed species |  | Q1 | Q2 | Q3 | Q4 | Q5 | Total | Frequency | Density | Relative frequency | Relative density | Importance value | Abundance | Dominance | Relative dominance |
| --- | --- | --- | --- | --- | --- | --- | --- | --- | --- | --- | --- | --- | --- | --- | --- |
| 1 | *Ageratum conizoides* Linn. | 22 | 23 | 30 | 15 | 17 | 107 | 5 | 85.6 | 9.4 | 15.5 | 25.0 | 12.5 | 1712.0 | 15.1 |
| 2 | *Bidens pilosa* Linn. | 48 | 30 | 26 | 18 | 32 | 154 | 5 | 123.2 | 9.4 | 22.4 | 31.8 | 15.9 | 2464.0 | 21.8 |
| 3 | *Borreria ocymoides* (Burm. f.) DC. | 8 | 16 | 18 | 20 | 20 | 82 | 5 | 65.6 | 9.4 | 11.9 | 21.3 | 10.7 | 1312.0 | 11.6 |
| 4 | *Commelina benghalensis* L. | 16 | 35 | 17 | 16 | 30 | 114 | 5 | 91.2 | 9.4 | 16.6 | 26.0 | 13.0 | 1824.0 | 16.1 |
| 5 | *Euphorbia hirta* Linn. | 4 | 3 | 12 | 2 | 4 | 25 | 5 | 20.0 | 9.4 | 3.6 | 13.1 | 6.5 | 400.0 | 3.5 |
| 6 | *Mariscus alternifolius* Vahl. (=M. *umbellatus* Vahl.) | 12 | 15 | 7 | 8 | 3 | 45 | 5 | 36.0 | 9.4 | 6.5 | 16.0 | 8.0 | 720.0 | 6.4 |
| 7 | *Peperomia pellucida* (L.) H. B. & K. | 2 | 0 | 6 | 4 | 1 | 13 | 4 | 13.0 | 7.5 | 2.4 | 9.9 | 5.0 | 325.0 | 2.9 |
| 8 | *Digitaria horizontalis* Willd. | 1 | 0 | 1 | 0 | 0 | 2 | 2 | 4.0 | 3.8 | 0.7 | 4.5 | 2.3 | 200.0 | 1.8 |
| 9 | *Setaria barbata* (Lam.) Kunth | 18 | 32 | 14 | 25 | 20 | 109 | 5 | 87.2 | 9.4 | 15.8 | 25.3 | 12.6 | 1744.0 | 15.4 |
| 10 | *Spigelia anthelmia* Linn. | 5 | 1 | 1 | 0 | 2 | 9 | 4 | 9.0 | 7.5 | 1.6 | 9.2 | 4.6 | 225.0 | 2.0 |
| 11 | *Talinum triangulare* (Jacq.) Willd. | 2 | 3 | 1 | 1 | 0 | 7 | 4 | 7.0 | 7.5 | 1.3 | 8.8 | 4.4 | 175.0 | 1.5 |
| 12 | *Tridax procumbens* Linn. | 5 | 2 | 0 | 1 | 1 | 9 | 4 | 9.0 | 7.5 | 1.6 | 9.2 | 4.6 | 225.0 | 2.0 |

WAI: weeks after intercropping, Q 1-5: Quadrat 1-5.

**Table 12.** Weed flora and species parameters for control plot after harvesting

| Weed species |  | Q1 | Q2 | Q3 | Q4 | Q5 | Total | Frequency | Density | Relative frequency | Relative density | Importance value | Abundance | Dominance | Relative dominance |
| --- | --- | --- | --- | --- | --- | --- | --- | --- | --- | --- | --- | --- | --- | --- | --- |
| 1 | *Bidens pilosa* Linn. | 0 | 4 | 5 | 1 | 1 | 11 | 4 | 11.0 | 8.9 | 1.6 | 10.5 | 5.2 | 275.0 | 1.8 |
| 2 | *Borreria ocymoides* (Burm. f.) DC. | 174 | 22 | 62 | 38 | 56 | 422 | 5 | 337.6 | 11.1 | 48.4 | 59.5 | 29.8 | 6752.0 | 44.0 |
| 3 | *Brachiaria deflexa* (Schumach.) C.E. Hubbard ex Robyns | 14 | 6 | 19 | 6 | 11 | 56 | 5 | 48.8 | 11.1 | 6.4 | 17.5 | 8.8 | 896.0 | 5.8 |
| 4 | *Brachiaria lata*(Schumach) C.E. Hubbard | 13 | 9 | 11 | 19 | 15 | 67 | 5 | 53.6 | 11.1 | 7.7 | 13.8 | 9.4 | 1072.0 | 7.0 |
| 5 | *Commelina benghalensis* L. | 17 | 27 | 18 | 29 | 27 | 118 | 5 | 94.4 | 11.1 | 13.5 | 24.6 | 12.3 | 1888.0 | 12.3 |
| 6 | *Digitaria horizontalis* Willd. | 0 | 6 | 1 | 5 | 0 | 12 | 3 | 16.0 | 6.7 | 2.3 | 9.0 | 4.5 | 533.3 | 3.5 |
| 7 | *Euphorbia heterophylla* Linn. | 0 | 0 | 2 | 0 | 0 | 2 | 1 | 8.0 | 2.2 | 1.1 | 3.4 | 1.7 | 800.0 | 5.2 |
| 8 | *Hyptis lanceolata* Poir. | 3 | 0 | 2 | 2 | 7 | 14 | 4 | 14.0 | 8.9 | 2.0 | 10.9 | 5.4 | 350.0 | 2.3 |
| 9 | *Mariscus alternifolius* Vahl. (=M. *umbellatus* Vahl.) | 14 | 11 | 5 | 0 | 3 | 33 | 4 | 23.0 | 8.9 | 4.7 | 13.6 | 6.8 | 825.0 | 5.4 |
| 10 | *Sida acuta* Burm. f. | 0 | 1 | 0 | 1 | 1 | 3 | 3 | 4.0 | 6.7 | 0.6 | 7.2 | 3.6 | 133.3 | 0.9 |
| 11 | *Sporobolous pyramidalis* P. Beauv. | 32 | 10 | 10 | 18 | 13 | 83 | 5 | 66.4 | 11.1 | 9.5 | 20.6 | 10.3 | 1328.0 | 8.7 |
| 12 | *Talinum triangulare* (Jacq.) Willd. | 2 | 6 | 3 | 0 | 0 | 11 | 3 | 14.7 | 6.7 | 2.1 | 8.8 | 4.4 | 488.9 | 3.2 |

WAI: weeks after intercropping, Q 1-5: Quadrat 1-5.

In tomato 01665-oil palm intercrop plot, at 3 WAI, *Borreria ocymoides* and *Sida acuta* recorded the highest and lowest values of relative importance values (RIV) of 46.3 and 6.6 respectively. At 6 WAI, *Peperomia pellucida* recorded the highest the RIV value (55.9), and *Sida acuta* and *Commelina benghalensis* both recorded the lowest (5.3). After harvesting, *Ageratum conizoides* and *Physalis angulata* both recorded the highest and lowest RIV values (45.1 and 7.4) respectively.

In tomato (NG/AA/SEP/09/053)-intercrop plot, at 3 WAI, *Ageratum conizoides* and *Borreria ocymoides* recorded the highest RIV (26.1); the lowest RIV was recorded by *Euphorbia hirta* (3.3). At 6 WAI, *Setaria barbata* recorded the highest RIV, while *Mariscus alternifolius* and *Talinum triangulare* both recorded the lowest RIV (7.9). After harvesting, *Brachiaria deflexa* and *Euphorbia hirta* recorded the highest and lowest RIV (46.3 and 6.0) respectively.

In eggplant (NGB 01737)-intercrop plot, at 3 WAI, *Synedrella nodiflora* and *Bidens pilosa* recorded the highest and lowest RIV values (42.7 and 4.7). At 6 WAI, the highest and lowest RIV values were recorded by *Ageratum conizoides* and *Brachiaria lata* (38.4 and 3.4) respectively. After harvesting, *Borreria ocymoides* and *Brachiaria lata* recorded the highest and the lowest RIV values (34.2 and 6.6) respectively.

In the control plot, at 3 WAI, *Borreria ocymoides* recorded the highest RIV (28.0) while *Commelina benghalensis* recorded the lowest value (6.2). At 6 WAI, *Bidens pilosa* and *Digitaria horizontalis* recorded the highest and lowest RIV values (31.8 and 4.5) respectively. After harvesting, *Borreria ocymoides* and *Sida acuta* recorded the highest and lowest values of RIV (59.5 and 7.2) respectively.

Of all the weed species that recorded highest RIV values across the plots and periods during which the weed sampling was carried out, *Borreria ocymoides* recorded the highest occurrence, by recording highest values of RIV five times. This was followed by *Ageratum conizoides* which recorded the highest occurrence, by recording highest values of RIV three times. On the other hand, of all the weed species that recorded lowest RIV values across the plots and periods during which the weed sampling was carried out, *Sida acuta* recorded the highest occurrence, by recording lowest values of RIV three times. *Commelina benghalensis*, *Euphorbia hirta* and *Brachiaria lata*, each recorded lowest RIV values across the plots and periods during which the weed sampling was carried out twice.

### Effect of intercropping distance on weed species abundance and richness in juvenile oil palm-tomato (NGB 01665) plot

The weed species richness at 3 and 6 WAI was highest in the control plot (11.80 and 10.80) respectively, however, at the end of harvesting, sub-plots at 2 and 3 m from the juvenile oil palm row recorded the highest weed species richness (10.00) (Table 13). Weed species abundance at 3 WAI was highest at 3 m from the juvenile oil palm (111.60), however; it was not significantly different from the control plot and other intercropping distances at (P<0.05) level of significance. At 6 WAI and after harvesting, the highest weed species abundance was recorded by the control plot (125.80 and 162.60), and were significantly different from the intercropping distances at (P<0.05) level of significance.

**Table 13.** Effect of intercropping distance on weed species abundance and richness in juvenile oil palm – tomato (NGB 01665) plot

| Intercropping distance | Weed species richness |  |  | Weed species abundance |  |  |
| --- | --- | --- | --- | --- | --- | --- |
|  | 3 WAI | 6 WAI | End of harvest | 3 WAI | 6 WAI | End of harvest |
| 1 m | 10.200<sup>b</sup> | 9.800<sup>ab</sup> | 7.800<sup>b</sup> | 88.400<sup>a</sup> | 89.400<sup>b</sup> | 83.400<sup>b</sup> |
| 2 m | 10.800<sup>ab</sup> | 8.600<sup>b</sup> | 10.000<sup>a</sup> | 104.60<sup>a</sup> | 93.200<sup>b</sup> | 98.800<sup>b</sup> |
| 3 m | 10.000<sup>b</sup> | 8.400<sup>b</sup> | 10.000<sup>a</sup> | 111.600<sup>a</sup> | 98.400<sup>b</sup> | 117.20<sup>b</sup> |
| Control | 11.800<sup>a</sup> | 10.800<sup>a</sup> | 9.000<sup>ab</sup> | 103.200<sup>a</sup> | 125.800<sup>a</sup> | 162.600<sup>a</sup> |

Means with the same letter(s) along the same column are not significantly different at (P<0.05)
level of significance.

### Effect of intercropping distance on weed species abundance and richness in juvenile oil palm-tomato (NG/AA/SEP/09/053) plot

The highest weed species richness value at 3 WAI was recorded at 1 m from the juvenile oil palm sub-plot (12.00), followed closely by the control plot (11.80). the lowest weed species richness value was recorded at 2 m from the juvenile oil palm sub-plot. At 6 WAI and at the end of harvesting, the control plot recorded the highest values of weed species richness (10.80 and 9.00) respectively. Weed species abundance at 3 WAI was highest in the control plot (103.20), and was not significantly different from the 3 m from the juvenile oil palm sub-plot (79.20) (P<0.05). At 6 WAI and at the end of harvest, the control plot similarly recorded the highest value of weed species abundance (125.80 and 162.60) respectively. That shows a progressive increase in the weed species abundance in the control plot from 3 WAI through to the end of harvest. The lowest values of weed species abundance at 6 WAI and at the end of harvest were recorded at 2 and 1 m from the juvenile oil palm sub-plot (50.40 and 55.80) respectively (Table 14).

**Table 14.** Effects of intercropping distance on weed species abundance and richness in juvenile oil palm – tomato (NG/AA/SEP/09/053) plot

| Intercropping distance | Weed species richness |  |  | Weed species abundance |  |  |
| --- | --- | --- | --- | --- | --- | --- |
|  | 3 WAI | 6 WAI | End of harvest | 3 WAI | 6 WAI | End of harvest |
| 1 m | 12.000<sup>a</sup> | 7.800<sup>b</sup> | 7.800<sup>a</sup> | 65.400<sup>b</sup> | 56.000<sup>c</sup> | 55.800<sup>b</sup> |
| 2 m | 9.200<sup>b</sup> | 7.800<sup>b</sup> | 8.600<sup>a</sup> | 57.200<sup>b</sup> | 50.400<sup>c</sup> | 77.200<sup>b</sup> |
| 3 m | 10.200<sup>ab</sup> | 9.600<sup>a</sup> | 8.800<sup>a</sup> | 79.200<sup>ab</sup> | 81.600<sup>b</sup> | 94.400<sup>b</sup> |
| Control | 11.800<sup>a</sup> | 10.800<sup>a</sup> | 9.000<sup>a</sup> | 103.200<sup>a</sup> | 125.800<sup>a</sup> | 162.600<sup>a</sup> |

Means with the same letter(s) along the same column are not significantly different at (P<0.05)
level of significance.

### Effect of intercropping distance on weed species abundance and richness in juvenile oil palm-eggplant (NGB 01737) plot

At 3 WAI, highest weed species richness was recorded at 1 m from the juvenile oil palm sub-plot (13.60), and was significantly different from the control plot and other intercropping distances at (P<0.05) level of significance. The control plot recorded the highest weed species richness at 6 WAI (10.80), while the lowest value was recorded at 1 m from the juvenile oil palm sub-plot (7.80). At the end of harvest, the highest weed species richness was recorded by both the control plot and at 2 m from the juvenile oil palm sub-plot, however, they were not significantly different from other intercropping distances at (P<0.05) level of significance. Highest weed species abundance value at 3 WAI was recorded at 3 m from the juvenile oil palm sub-plot (122.00). At 6 WAI, the highest and the lowest weed species abundance values were recorded by the control plot and at 1 m from the juvenile oil palm sub-plot (125.80 and 63.20) respectively. Similarly, at the end of harvest, the control plot recorded the highest weed species abundance (162.60), while the lowest was recorded by those at 1 m from the juvenile oil palm sub-plot (76.00) as shown in Table 15.

**Table 15.** Effects of intercropping distance on weed species abundance and richness in juvenile oil palm – eggplant (NGB 01737) plot

| Intercropping distance | Weed species richness |  |  | Weed species abundance |  |  |
| --- | --- | --- | --- | --- | --- | --- |
|  | 3 WAI | 6 WAI | End of harvest | 3 WAI | 6 WAI | End of harvest |
| 1 m | 13.600<sup>a</sup> | 7.800<sup>c</sup> | 8.600<sup>a</sup> | 82.800<sup>b</sup> | 63.200<sup>d</sup> | 76.000<sup>b</sup> |
| 2 m | 11.200<sup>b</sup> | 8.800<sup>bc</sup> | 9.000<sup>a</sup> | 79.600<sup>b</sup> | 82.800<sup>c</sup> | 87.20<sup>b</sup> |
| 3 m | 12.400<sup>b</sup> | 9.600<sup>ab</sup> | 8.600<sup>a</sup> | 122.000<sup>a</sup> | 97.000<sup>b</sup> | 101.800<sup>b</sup> |
| Control | 11.800<sup>b</sup> | 10.800<sup>a</sup> | 9.000<sup>a</sup> | 103.200<sup>ab</sup> | 125.800<sup>a</sup> | 162.600<sup>a</sup> |

Means with the same letter(s) along the same column are not significantly different at (P<0.05) level of significance.

### Simpson’s Diversity indices of juvenile oil palm-vegetable intercrops

At 3 WAI, the intercrop plots with highest and lowest weed species diversity values as informed by the values of Simpson Diversity Index (D) are juvenile oil palm-tomato (NG/AA/SEP/09/053) plot and the control plot (0.186 and 0.242) respectively. At 6 WAI, the highest diversity was recorded at the control plot with Simpson’s Diversity Index of 0.123, while the lowest diversity was recorded at juvenile oil palm-tomato (NGB 01665) plot, with Simpson’s Diversity Index of 0.266. At the end of harvest, the juvenile oil palm-eggplant (NGB 01737) plot recorded the highest Simpson’s Diversity Index of 0.156 (Table 16).

**Table 16.** Simpson’s Diversity indices of juvenile oil palm-vegetable intercrops at 3 WAI, 6 WAI and after harvesting

| Simpson's diversity | 3 WAI |  |  | 6 WAI |  |  | End of harvest |  |  |
| --- | --- | --- | --- | --- | --- | --- | --- | --- | --- |
|  | D | 1-D | <sup>1</sup>/D | D | 1-D | <sup>1</sup>/D | D | 1-D | <sup>1</sup>/D |
| Plot A | 0.198 | 0.802 | 5.051 | 0.266 | 0.734 | 3.759 | 0.200 | 0.800 | 5.000 |
| Plot B | 0.186 | 0.814 | 5.376 | 0.173 | 0.827 | 5.780 | 0.201 | 0.799 | 4.975 |
| Plot C | 0.195 | 0.805 | 5.128 | 0.213 | 0.787 | 4.695 | 0.156 | 0.844 | 6.410 |
| Plot D | 0.242 | 0.758 | 4.132 | 0.123 | 0.877 | 8.130 | 0.300 | 0.700 | 3.333 |

Plot A – D: juvenile oil palm – tomato (NGB 01665), juvenile oil palm – tomato (NG/AA/SEP/09/053), juvenile oil palm – eggplant (NGB 01737) and control plot.
D: Simpson's diversity index, 1 – D: Simpson's index of diversity and <sup>1</sup>/D: Simpson's reciprocal index of diversity.

## Discussion

The reduction observed in number of weed species at 3 through to 6 WAI and after harvesting of fruits may be due to the shading effect of the intercropped juvenile oil palm and also the canopy shade cast by the fruit vegetables. In addition, the effect of tilling of the soil during land preparation may result in reduction of number of weed species at this period. Soil tillage could cause weed seed to be buried deep in the soil where factors such as unsuitable temperature at that depth could prevent their germination. This finding agrees with the submission of Baker *et al.* (2018), who studied weed species composition and density under conservative agriculture and reported that weed seeds accumulate in the top soil in conservative agriculture-in which little or reduced tillage was practiced. They suggested that it could accelerate weed seed germination when conditions are favourable. Similarly, this observation corroborates an earlier study by Weber *et al.* (2017), who worked on weed control using conventional and reduced tillage, no-tillage and cover crops in organic soybean. They reported that weed seeds abound in the top soil due to the absence of tillage, and these seeds could easily germinate when conditions are favourable. This observation is also in tandem with the submission of Benvenuti *et al.* (2001) who carried out a research on emergence of seedlings from buried weed seeds with increasing soil depth. They ascribed prompt weed growth at the top soil to the availability of favourable germination conditions at that soil layer. Besides, they submitted that in non-moisture-limiting conditions, germination stimulus is generally higher near the soil surface which is light-rich with diurnal temperature fluctuations.

Considerably high numbers of broadleaf weeds were reported in most of the intercropped plots in comparison to grasses and this observation may be due to abundance of seeds and propagules of these species in soil seedbank of shaded or partly shaded soil in tropical rainforest zone-where the experimental juvenile oil palm plantation used in this study is located. Furthermore, this finding could also be explained by the ecological conditions present in shaded environment with regard to light availability and moisture level. The canopy cover offered by the juvenile oil palm could lead to reduced light availability for grasses, thereby promoting the growth of broadleaf weeds. The abundance of broadleaf weeds observed during this study is in consonance with as earlier study by Olorunmaiye *et al.* (2011) who investigated the weed species composition of citrus-based cropping systems. They reported an elevated concentration of broad leaf weeds beneath the canopies of adult citrus trees. Similar submission was made by Kumar *et al.* (2010) who studied the performance of ginger in intercrop and in sole cropping. They reported higher density of broadleaf weeds in intercrop than observed in sole cropping.

Of the broadleaf weeds recorded during this study, members of the families Asteraceae and Euphorbiaceae were observed to have higher numbers of members at 3 and 6 weeks after intercropping and after harvesting. This observation could be attributed to the lifecycle, resources demands and seed dispersal mechanism of these families of weeds. This observation is in accordance with a prior report by Olorunmaiye *et al*. (2011) who suggested high colonizing power of these two families, readily brought about by the high fruit production and the efficient dispersal of fruits and seeds. This is further corroborated by a report by Oluwatobi and Olorunmaiye (2014) who probed the weed species distribution of vegetable and arable crops intercropped with juvenile oil palm. They identified towering light requirement, aggressive growth, short life cycle, and large seed production with potent volatile seed dispersal mechanism as attributes that may be responsible for the high relative weed density observed in members of Asteraceae and Euphorbiaceae.

Some weed species such as *Mariscus alternifolius*, *Borreria ocymoides*, *Ageratum conizoides and Euphorbia heterophylla* were found to be present in most of the plots sampled. This may be because these weeds are predominant in juvenile oil palm field of the study area, constituting larger proportion of weed seeds in the soil seedbank. This finding is in line with previous study by Yakubu *et al*. (2006) who reported site specificity in both crops and weeds. Closely related finding was also opined by Karaye *et al*. (2007), who reported that some weeds and crops are site specific, whereas others will thrive over an extensive range of habitat.

Weed species abundance and richness were found to be higher in non-intercropped plot when compared to the intercropped plot (control plot). This could be ascribed to the difference in the biotic, abiotic or ecological conditions provided by the intercropped and non-intercropped plots. This observation is complemented by the finding of an earlier report by Sosnoskie *et al.* (2006) who investigated the weed seedbank community composition and suggested that monoculture promotes weed species abundance and richness.

The control plots recorded lower weed species diversity as shown by the Simpson’s Index values at 3 WAI and at the end of harvesting. This observation could be explained by the absence of canopy cover that would have offered shade as found in the intercropped plot, by the juvenile oil palm. This absence of shade could promote weed diversity, as observed in the control plot. This observation is in harmony with an earlier report by Baker *et al.* (2018), who evaluated the effect of cropping system on weed density and diversity. They put forward that higher weed diversity was recorded in non-intercrop plot as compared to intercrop plot. They asserted that intercropping practice helps to effectively control weeds, ascribing it to the deleterious effects it has on weed growth and reproduction. More so, this observation also affirms earlier report by Obadoni *et al.* (2009) who reported that shade upshot from trees did not promote undergrowth rejuvenation or establishment of weed.

Additionally, this observation is also supported by a study by Mhlanga *et al.* (2016) who reported reduced weed diversity through pronounced shading of weeds. Further, a study by Rahimi *et al*. (2019) who investigated the influence of intercropping and weed management practices on weed parameters, is in unison with the finding of this work. They recorded higher diversity of weed species in intercropped plot than in sole cropping.

## Conclusions

There was progressive reduction in the number of weed species composition from 3 WAI through to after harvesting. Intercropping fruit vegetables with juvenile oil palm resulted in more abundance of broadleaf weed species than grass species. The most predominant broadleaf weed species belong to families Asteraceae and Euphorbiaceae. *Mariscus alternifolius*, *Borreria ocymoides*, *Ageratum conizoides and Euphorbia heterophylla* were found to be present in most of the plots sampled. Higher weed species abundance and richness were observed in the control plot where sole fruit vegetable crops were cultivated. On the other hand, the intercropping of fruit vegetables with juvenile oil palm promoted higher diversity of weed species than recorded in the control plot. Intercropping of fruit vegetables within the alley of juvenile oil palm should be encouraged among farmers, especially at the earlier years before canopy closure.

## Acknowledgements

This research received no specific grant from any funding agency in the public, commercial, or not-for-profit sectors.

